# Transient Magnetic Resonance Elastography: a method to measure the mechanics of the active heart

**DOI:** 10.64898/2026.03.02.708995

**Authors:** Marco Barbero Mota, Giacomo Annio, Guillaume Rucher, Jordi Martorell

**Affiliations:** U1148, Laboratory for Vascular Translational Science, INSERM, Paris, France; Fédération de Recherche en Imagerie Multimodalité - UMS34 Claude Bernard INSERM, Paris, France; Department of Chemical Engineering and Material Sciences, IQS School of Engineering, Barcelona, Spain

## Abstract

Myocaridum biomechanics are a biomarker for multiple cardiac pathologies. However the rapid and complex heart motion hampers accurate measurements of the tissue stiffness. Current *in vivo* methods for the evaluation of myocardium mechanical health are either highly invasive or can only provide with a global surrogate of heart function as they suffer from poor spatiotemporal resolution. We propose a new *in vivo* technique, transient magnetic resonance elastography (tMRE), to assess the dynamic cardiac biomechanics. tMRE is able to quantify local shear wave speed as a proxy for myocardial stiffness at user-defined times within the cardiac cycle. We report proof-of-concept results where we probe the septum of 4 different healthy rat specimens at 3 physiologically distinct cardiac phases. We provide with apparent speed measurements for early systole, mid-late systole and early diastole that match the expected values from the cardiac cycle physiological mechanics. We correct for non-negligible geometrical biases using literature results and report true stiffness values where possible. Finally, we validate tMRE in phantom experiments.

## Introduction

The heart is a contractile mechanical pump whose biomechanics are key to understanding its healthy functioning, as well as to diagnose and predict any present or future malfunctioning^1^. Heart function relies both on the active/contractile and the passive/relaxation phases also known as systole and diastole.

The mechanics involved in cardiac contraction have been thoroughly studied through various attempts to measure the stiffness of the left ventricle (LV) tissue. *Ex vivo*^2^ and computational^3^ methods make assumptions regarding the myocardium *in vivo* conditions and thus do not account for all the factors that play a role in determining the mechanics of the heart *in situ*. Measurements from these incomplete tissue models need to be carefully assessed when used for *in vivo* and clinical applications.

Non-invasive *in vivo* methods to measure systolic cardiac mechanics *in situ* have also been proposed, mainly in the imaging space. Cardiac magnetic resonance imaging, ultrasound, and computer tomography are the three families of imaging methods within which multiple variants have been proposed []. In general, these tools aim to measure changes in ventricular volume and use this information to provide an estimate of tissue stiffness at the organ level^1^. However, this is insufficient for clinical purposes that require local metrics. Focal and minor, but crucial, deviations from normal cardiac function may pass unnoticed if a tool is not sensitive to biomechanical changes at the sub-organ level. These slight abnormalities usually characterize CVDs in the early stages when treatments are most effective. [][]

Although most heart failures (HF) develop under systolic symptomatology, some subtypes have their root cause in abnormal relaxation during diastole. These conditions are known as heart failure with preserved ejection fraction (HFpEF) in which the heart is still able to fully eject the blood pool but is unable to completely fill during relaxation. Stroke volume is therefore reduced causing a dwindled cardiac output^4,5^. These conditions affect more than half of patients with HF^6^ and more than 70% of the elderly population (>65 years of age)^7^. State-of-the-art *in vivo* methods for assessing *in situ* diastolic heart mechanics face significant limitations. Most methods struggle to separate passive and active mechanics due to limited temporal resolution, lack direct access to the myocardium, and often rely on indirect measures such as blood flow patterns or strain tracking that do not directly measure local myocardial stiffness^8^. Among them, echocardiography is still the gold standard technique used in the clinic. However, it can only provide an average surrogate estimate of LV stiffness that conflates various factors and may hide abnormalities in myocardial stiffness^9^.

Despite the plethora of available methods, the current gold standard for assessing *in situ* LV cardiac performance, both during systole and diastole, is still the relationship between ventricular pressure and volume (P-V diagram)^10,11^. It is obtained by catheterization, usually aided by an imaging technique. It provides an accurate measure of overall LV pumping performance^1^. The P-V diagram approach has two main disadvantages. First, it is highly invasive; thus its feasibility in clinical practice is reduced to critical cases that exclude its use for routine cardiac monitoring or with high risk patients. Second, it lacks spatial resolution as it can only provide a single measurement of the ventricular volume-pressure ratio that summarizes cardiac function at a specific timepoint. These global point estimates may hide local and/or punctual abnormalities discriminatory of the root cause(s) of different HFs such as myocardial fibrosis or cardiac wall infarction. Such conditions may affect distinct areas of heart tissue and may change myocardical biomechanics following different temporal patterns across the cardiac cycle.

The above shortcomings in the state of the art tooling to measure cardiac mechanics call for non-invasive approaches that can directly measure local passive myocardial stiffness *in vivo* and *in situ*. Such tool must be equipped with high spatiotemporal resolution to be able to detect changes in myocardium stiffness both during the active contractile phase as well as during passive relaxation. Such technique would allow continuous patient monitoring and therefore aid clinicians in routine tasks for CVDs diagnosis, prognosis and in the assessment of CVDs therapy response.

Magnetic resonance elastography (MRE) is a phase-contrast magnetic resonance imaging (MRI) technique to measure the intrinsic mechanical properties of tissue^12^. Motion sensitive gradients capture tissue displacement using modulation in the phase of the MRI signal. Raw phase data is fed into an inversion algorithm that solves locally the wave equation in the region of interest. This algorithm generates a 3D elastogram, that is a map of the complex shear modulus (*G*^*^ = *G*^*′*^ + *iG*^*″*^, with *G*^*′*^ shear stiffness and *G*^*″*^ shear viscosity) of the tissue imaged.

Cardiac MRE is particularly challenging for several reasons. First, the dynamic nature of the heart makes data acquisition prone to motion artifact and requires high temporal resolution to be able to distinguish shear waves propagation through the myocardium at different cardiac stages. Second, the thin-wall geometry of the heart gives origin to a non-negligible wave-guidance effect. Under this phenomena, observed shear wave phase velocity is frequency-dependent and thus does not reflect the true material properties without correct debiasing. Clinical applications require a range of operational frequencies (∼ 100-500 Hz) in which this bias may be severe precluding the direct translation of apparent speed into true shear modulus^13–15^.

Despite these challenges, prior work demonstrated the feasibility of measuring shear wave speed *in vivo* in mouse models^16^, using an acoustic driver delivering 400 Hz steady state waves. Tissue displacement was measured using fractional encoding MRE. However, the proposed approach has several limitations. First, potential bias in shear wave speed values due to sensitivity to the longitudinal waves in steady state. Secondly, stationary MRE in thin structures is especially susceptible to waveguide effects which the authors did not account for the in their inversion algorithm and resulting elastogram. On this matter, Smith et al^17^ validated the use of MRE to distinguish between soft, intermediate and stiff biventricular rat heart phantoms. They found that the absolute values had been significantly impacted by the geometrical biases specially when the shear waves wavelength was longer than half the size of the heart.

Our work presents a novel transient cardiac MRE (tMRE) technique that aims to approach the problem of non-invasive *in vivo* cardiac stiffness measurement from a different angle than conventional cardiac MRE. We build on the work by Troelstra et al^15^ who instead of using steady state MRE and inversion of the raw phase data utilized intrinsic shear wave speed as a proxy for stiffness.

Cardiac tMRE allows for direct measurements of shear wave velocity within myocardium tissue at high spatial and temporal resolution. It also enables to probe the myocardium at any timepoint of interest throughout the cardiac cycle through the usage of external mechanical excitation.

As a proof of concept, we quantify apparent shear wave phase velocity for three arbitrary distinct cardiac phases, early systole (ES), mid-late systole (MS), and early diastole (ED), in the interventricular septum of rat subjects. We leverage a previously published thin plate bias correction model to correct for geometrical effects that obscure the true material properties. Finally, we delineate the limitations of this technique and future work needed to better remove these geometrical biases from shear wave velocity measurements.

## Methods

### tMRE data acquisition (DAQ)

tMRE was performed on 4 male 10-12 weeks old wistar white rats (Janvier Labs, Le Genest-Saint-Isle, France). The animals were anesthetized with 3% gaseous isoflurane-air mixture and placed on an in-house custom made 3D printed set-up. This system consists of a half-cylindrical bed with a semi-rigid polyethylene axis connected to a piston through a polyether ether ketone (PEEK) lamella on one end, and to a linear electrodynamic motor on the opposite side, at the back of the MRI bore. The layout of the custom bed is such that the piston sticks out through a hole on the flat surface where the animal lays and is in direct contact with the animal’s thoracic cavity right in front of the heart. The piston was designed to serve as holder for the MRI radiofrequency receiver coil seeking to maximize electromagnetical signal on the animal chest (figure 1). This study was in compliance with the French government rules of the French animal experimentation ethics Autorisation de Projet utilisant des Animaux à des Fins Scientifique.

**Figure 1.**
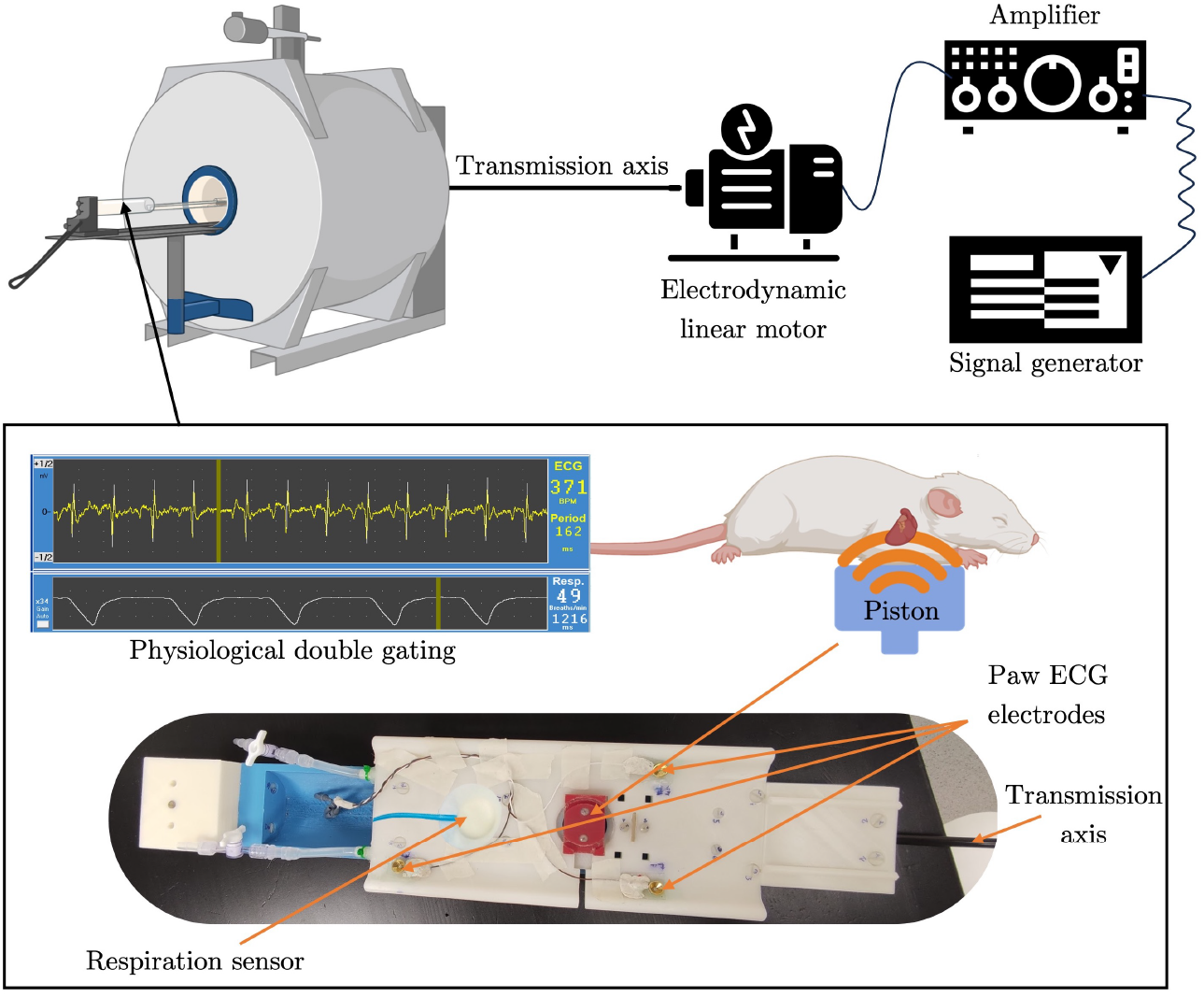
Transient MRE experimental setup. Our 3D printed bed (bottom) included a heating system for specimen comfort, nose cone for anesthesia (not pictured), both cardiac and breathing sensors for the sequence gating, and the mechanical system to transmit pulses from the motor into the animal chest: rigid transmission axis connected to a lamella onto which the piston is attached to transduce the vibrations.

Custom pulse signals were sent from a signal generator (Tektronix AFG3022C; Dual Channel arbitrary function generator 250MS/s 25Mhz) and amplified (75 VA Power Amplifier, Type 2718, Brüel & Kjær) before being inputted into an electrodynamic linear motor (Mini-shaker Type 4810, Brüel & Kjær) to generate the corresponding mechanical pulses. The motor longitudinal vibrations are transmitted via the transmission axis to a cantilever-like assembly which sustains the piston in contact with the thoracic cavity tissue (figure 1).

Due to the dynamic nature of the heart, we use a double gated sequence via surface ECG electrodes and a balloon-like respiratory sensor positioned on the bed just underneath the animals’ limbs ends and lower abdomen, respectively. The occurrence of a R-peak between end of exhalation and the start of inhalation triggers the tMRE sequence.

We used a 2D Gradient Echo Flash CINE sequence (TE=2.3 ms; TR=13ms; FOV 64 mm; matrix 128 × 128; 1 slice; resolution 1 × 1 × 5mm^3^) equipped with motion sensitive bipolar gradients (1ms length, 150 mT/m strength) to measure shear waves-induced phase perturbations. Axial planes were positioned intersecting the left and right ventricles at their maximum short heart axis radii at the mid section of the heart long axis. To minimize blood flow derived motion artifacts, we equipped the sequence with first order gradient moment nulling and a flow saturation prepulse positioned on the atria. All tMRE data acquisition was performed on a preclinical 7T MRI scanner (Biospec 70/20, Bruker, Germany). Standard coil tuning and matching (wobbling adjustment) were performed at the beginning of every experiment as well asllipsoidal volume shimming centered at the specimen’s heart was routinely done prior to every tMRE acquisition.

Each tMRE sequence consisted of 11 movie frames covering the whole cardiac cycle, acquired at R-peak delays between 13 to 146 ms temporally equidistant assuming an average cardiac cycle duration of 160 ms. 166.67 Hz full sine pulses starting at 15, 52 and 104 ms delays relative to the beginning of trigger R-peak were used to probe the septum during ES, MS and ED, respectively (figure S1).

To enhance the otherwise limited temporal resolution of one TR (13 ms) our tMRE approach requires of a user-defined resolution factor, *k*. We found 100 to be the optimal value for our experiments. Image acquisition was repeated *k* + 1 times per experiment. For the first *k* repetitions, the mechanical excitation was retarded by a cumulative delay of *TR/k* = 0.13ms, while for the last shot no mechanical excitation was sent into the specimen.

Image acquisition was locked to the R cycle for all 11 movie frames which enabled us to retrospectively re-order all 1100 snapshots to artificially increase our temporal resolution down to *TR/k* = 0.13ms. Figure 2 graphically summarizes the tMRE data acquisition process.

**Figure 2.**
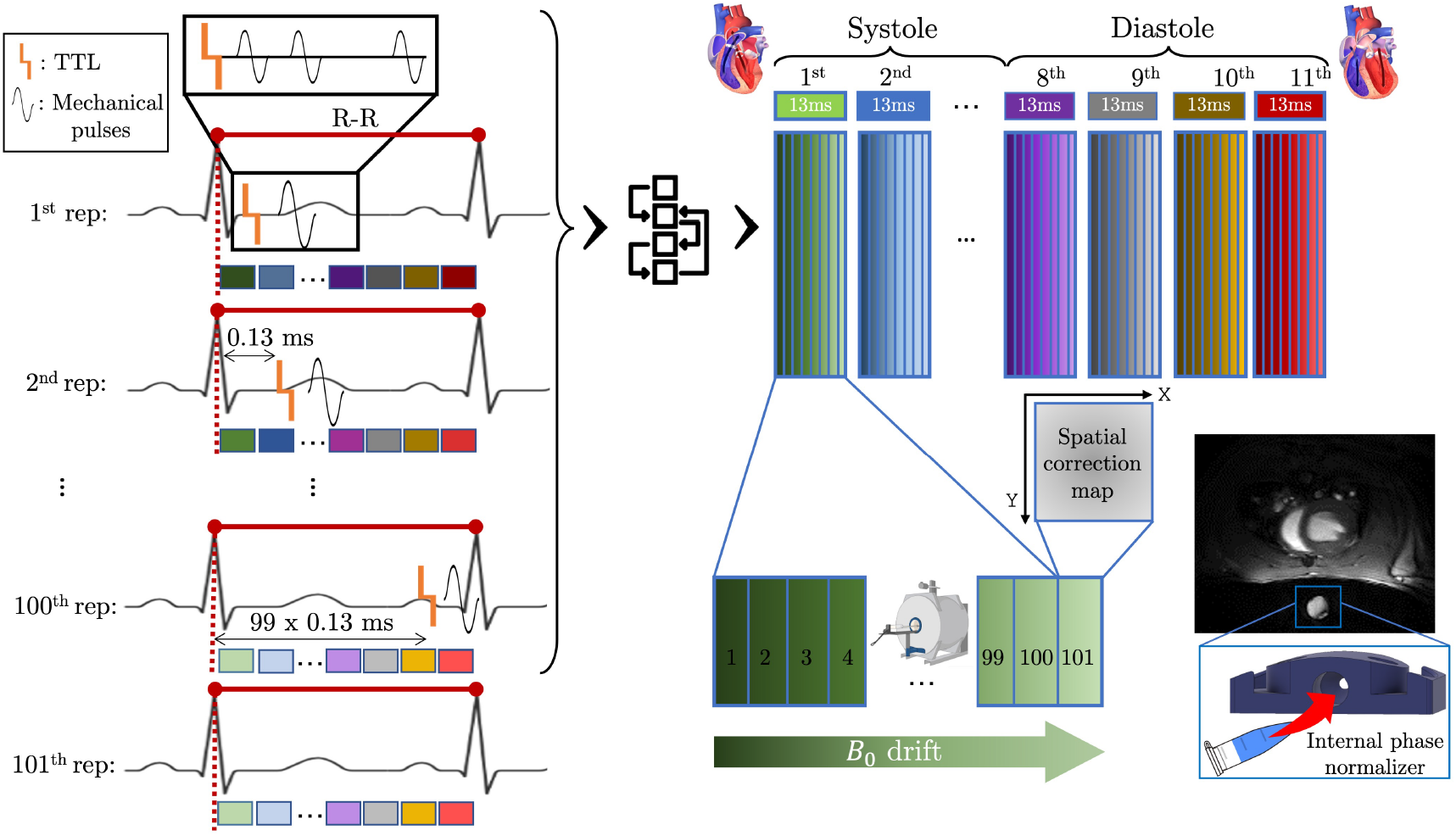
The imaging of 11 movie frames is repeated 100 times with a 0.13 ms cumulative delay in the mechanical excitation. An extra 101^th^ repetition of all movie frames is also acquired without mechanical excitation to serve as spatial correction maps. An ultrasound gel vial is embedded in the piston to be imaged within the FOV in all repetitions. The raw phase value of this vial is subtracted as phase baseline in each repetition (internal phase normalizer) to correct for any temporal drift. *TTL*: Transistor-Transistor Logic trigger from the MRI machine; *R-R*: ECG R-R interval; *B*_0_.: MRI static magnetic field.

In this study two complete independent tMRE data acquisitions were acquired for each of the 4 animals to demonstrate repeatability and reproducibility of our technique. Both tMRE repetitions were run sequentially without moving the specimen or the FOV.

After the second tMRE experiment the axial acquisition slices were also imaged at a higher in-plane spatial resolution (0.33 × 0.33 × 5 mm^3^) using a T1 fast low angle shot (FLASH) sequence. 11 anatomical acquisitions were acquired per each animal, one corresponding to each tMRE movie frame.

### Raw tMRE data preprocessing - temporal and spatial drift correction

Magnetic field heterogeneities were of special concern during tMRE imaging, both in the spatial and temporal dimensions. Phase shift artifacts can significantly bias the derived wave speed quantification from the full reordered phase movie data. On the one hand, the high 7T *B*_0_ static field imposes inhomogeneities even within the thin region of interest (i.e. interventricular septum). On the other hand, temporal drifts in the magnetic field throughout the data acquisition time can introduce phase offset gradients across repetitions for each movie frame. This is a phase artifact rooted in the retrospective reordering of the acquisitions whose effect was directly proportional to the duration of the experiment. Such duration is dependent on the physiological state of the animal given the double gating approach.

To reduce the effect of both of these artifacts in our resulting phase readout movies we perform a two-step normalization of all tMRE raw phase images (Figure 2):

- Spatial heterogeneity removal. At the end of each experiment we acquire an extra repetition to create spatial correction maps, one for each each movie frame (figure 2). This additional repetition encodes the remaining magnet intrinsic heterogeneities in the field of view (FOV).
- Temporal magnet drift correction. The piston of our custom bed set-up was engineered to host a propylene glycol ultrasound gel phantom in its base (figure 2). We use this unperturbed tissue-like media as an in-FOV internal reference for each acquisition. We subtract its phase value to the corresponding repetition phase image. This correction avoids the phase shift caused by *B*_0_ temporal drift that would otherwise occur within each movie frame after the reordering of the 100 repetitions.

### Waterfall diagram and speed quantification

To quantify apparent speed values of shear waves travelling through the interventricular septum we obtain what we call waterfall diagrams. We select the adjacent pixels that lie on a line running through the septum tissue (space dimension) and we extract their raw phase signal across movie frames (time dimension) (figure 3 A and C). We then unwrap the phase data and compile it into the waterfall diagram where the X-axis is the space dimension and the Y-axis is time (figure 3 B).

**Figure 3.**
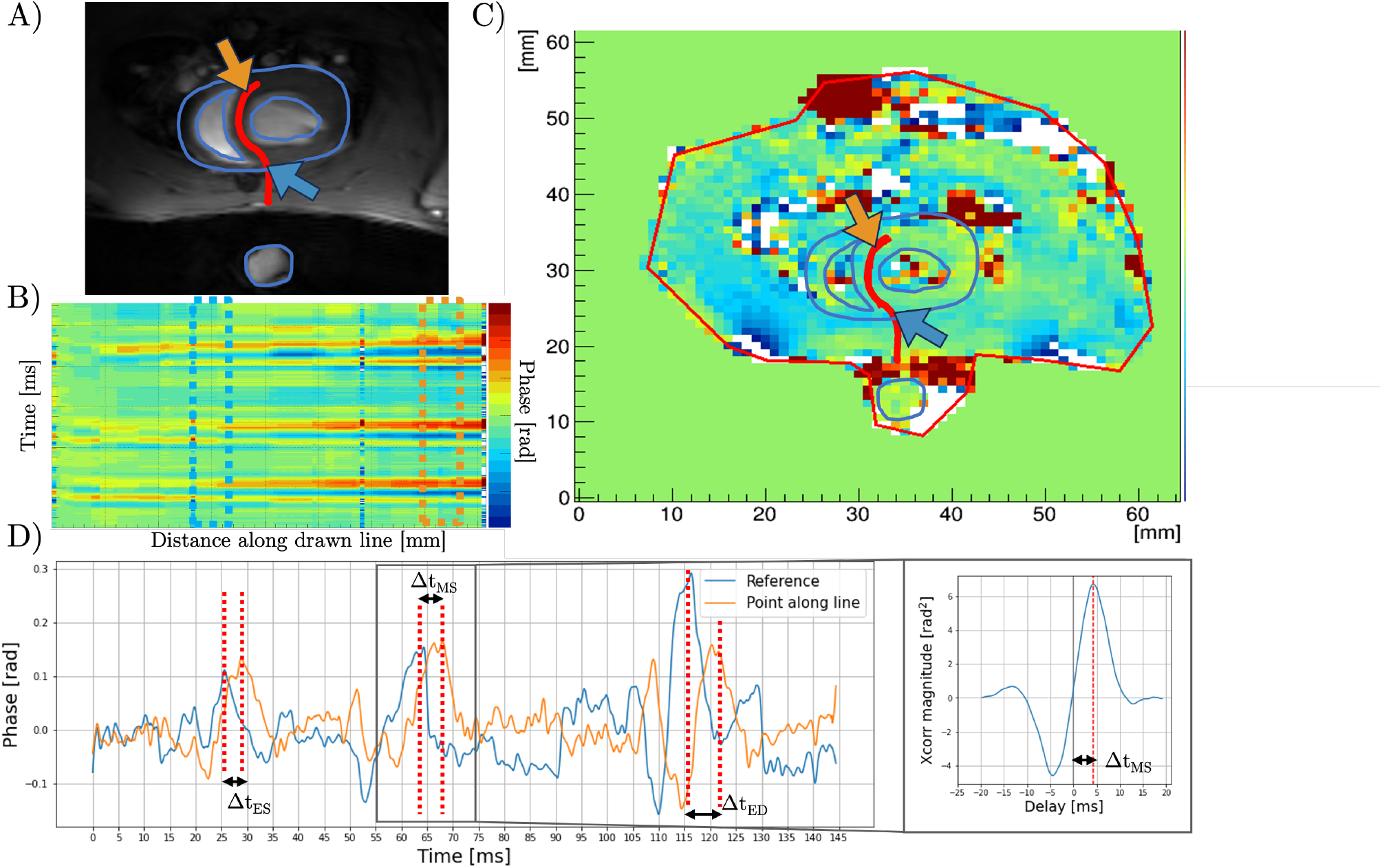
Blue and orange arrows, dashed highlighting boxes, and signals refer to the spatial locations used as reference and an arbitrary line point where shear wave speed is quantified. **A)** Drawn line superimposed to axial T1 anatomical image. **B)** Waterfall diagram displaying the propagation of the three shear waves through the space spanned by the drawn line. Note ED showing a steeper slope (slower wave) than ES and MS propagation which intuitively matches with the heart mechanical state. **C)** Unwrapped phase image for a particular movie frame repetition. **D)** Left: Unwrapped phase temporal signals for the reference and example line point. Red dashed lines indicate the measured delay for each cardiac phase. Larger for ED and smaller for ES and MS for the same distance traveled. Right: MS cross-correlation signal. Maximum peak finding gives the delay between line point and reference pulses.

Waterfall diagrams allow to identify the different wave modes induced into the septum based on their apparent speed. Our aim was to quantify the slower trailing edge of the shear waves from the waterfall diagram data as a proxy for shear modulus (*G*^*′*^)^15^. Shear waves phase perturbations leave oblique slopes in the waterfall diagram in contrast to longitudinal waves which travel much faster and display as horizontal lines (figure 3 B). Thus, to remove the unwanted phase disturbance we subtract the phase signal of the first pixel on the line to the rest of the diagram. Waterfall diagram generation was performed via an in-house C++ based graphical interface.

Each waterfall diagram data is then processed by an in-house script (Python 3.9.6) that runs the following pipeline to quantify apparent shear wave speed:

1. Smoothing: Each pixel phase signal is denoised via a Savitzky-Golay filter^18^ (window:11, order:2).
2. Reference point signal: We take the first pixel of the interventricular septum as reference point and isolate the externally induced perturbation phase signal. The rest of the temporal sequence is zero padded giving the *reference signal*.
3. Quantification: We loop over all the pixels on the region of interest (septum drawn line), starting a minimum of 0.5 mm away from the reference point. At each step we obtain the pixel temporal phase signal and perform the following steps:
  3.1 Signal preprocessing: We capture the full pulse of interest (ES,MS or ED) using the probing cardiac phase delay as starting point. We allow for a longer width of 35 ms to ensure that we catch the full sine pulse at every point along the line. We zero-pad the rest for a complete *line signal*.
  3.2 Delay estimation: We compute the cross-correlation function between the *reference* and the *line signal* and use maximum peak finding to estimate the delay that takes for the wave to travel between the reference pixel and the current position on the septum line. (figure 3 D).
  3.3 Speed computation: Knowing the delay and the distance between both the reference and the line point we can estimate shear wave velocity.
4. Filtering: We apply two filters to each experimental sample of speed estimations to remove unfeasible outliers:
  - We filter out any negative or *>* 20 m/s value. These samples most likely result from noisy SNR signals where maximum peaks correspond to noise spikes and not shear waves.
  - Low SNR *line signals* give unreliable cross-correlation signals. To minimize their effect in our estimations we assume a gaussian distribution of delay values 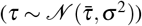 and disregard any outlier outside the conservative range 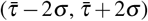.
5. Results: We compute mean and standard error on the mean for each experimental filtered speed sample.

We run this process three times for each experiment data, one per cardiac phase. The changing heart anatomy across the cardiac cycle requires tailoring the region of interest (drawn line) to obtain accurate waterfall diagrams.

For each animal, pooled statistics were calculated as weighted averages across both repetitions taking into account the amount of valid pixel speed values quantified within each experiment after outlier correction.

### Statistical modelling

We fitted a mixed effects model (MEM) to the mean speed data across animals and repetitions in order to analyze the differences in apparent speed between the three cardiac phases (ES, MS, ED). MEMs are normally used in scenarios of correlated data that can be clustered into subgroups, for example repeated measures for a certain group of individuals. MEMs have the property of separately estimating both random and systematic effects and consequently giving a more unbiased parameter estimation than vanilla linear regression^19^.

From the fitted MEM we obtained estimations for the random effects second-order statistical moments (variance and covariance). Such values allow for the computation of correlation coefficients (CCs) for ES and ED which can reveal patterns stemming from speed variability across cardiac phases. For details on model description, implementation and CCs computation see the Supplementary Section 2.1.

### Phantom validation

We validated tMRE apparent speed quantification pipeline on homogeneous ultrasound gel phantoms (10 × 10 × 2 cm^3^) of known stiffness. We performed the same experimental DAQ as with rat specimens by positioning the piston underneath the center of the phantom to approximate infinite medium conditions. We ran both the *in vivo* experiments tMRE sequence without double gating. We draw a region of interest line perpendicular to wave propagation to obtain a waterfall diagram from which shear wave speed is quantified as described above. In this case we obtain three estimations (ES, MS, and ED pulses) of the same quantity due to the static nature of the phantom. We translated speed into stiffness following:

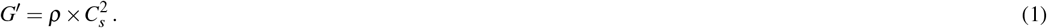

Our phantom media (ultrasound gel) has a known stiffness of *G*^*′*^∼ 1kPa. As an additional sanity check, we also verified such value via steady state MRE acquisition. Data acquisition was implemented as a multi-slice single spin echo sequence (TR = 900 ms; TE = 29 ms; FOV = 19.2 mm; matrix 64 × 64; 1 average; 8 wave phases; 9 slices; isotropic resolution 0.3 mm; acquisition time 23 min; vibration frequency = 200 Hz)^20^. We used the MRE inversion algorithm in Sinkus et al^21^ to reconstruct the phantom elastogram from which we sample regions of low non-linearity and at least 5 *µ*m displacement to obtain a stiffness estimation. We used the delta method^22^ (see Supplementary Section 2.2) to approximate *G*^*′*^_*phantom*_ standard errors.

### Geometrical bias correction

The geometry of the heart invalidates the infinite media assumption and imposes a non-negligible thin-plate bias effect on our apparent speed measurements. We used the experimental law proposed by Mo et al^23^ to correct for this effect. The authors proposed a non-linear relationship between true (*C*_*s*_) and apparent (*C*) speeds, parameterized by the frequency of vibration (*ν*) as well as the thickness of the thin structure (*H*):

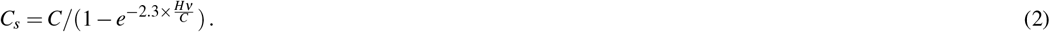

We estimated interventricular septum thickness (*H*) in each experiment through high resolution T1-FLASH anatomical images. We measured tissue thickness perpendicular to the tMRE quantification line 20 times along its length using ImageJ^24^ software. We obtained an estimator for *H* via the mean of each image sample. We then apply equation (2) to each animal’s mean apparent shear wave speed. The corrected speed value can then be transformed into shear modulus via equation (1). To obtain a valid measure of uncertainty for both *Cs* and *G*^*′*^ we approximate their standard errors using the delta method^22^ (see Supplementary Section 2.2)

## Results

### Apparent speed estimation and analysis

Quantified mean and error on the mean observed speed values are summarized in figure 4, grouped by animal for clarity. The same graph grouped by experiment (Rat-repetition pairs) can be found in the supplementary materials (figure S2).

**Figure 4.**
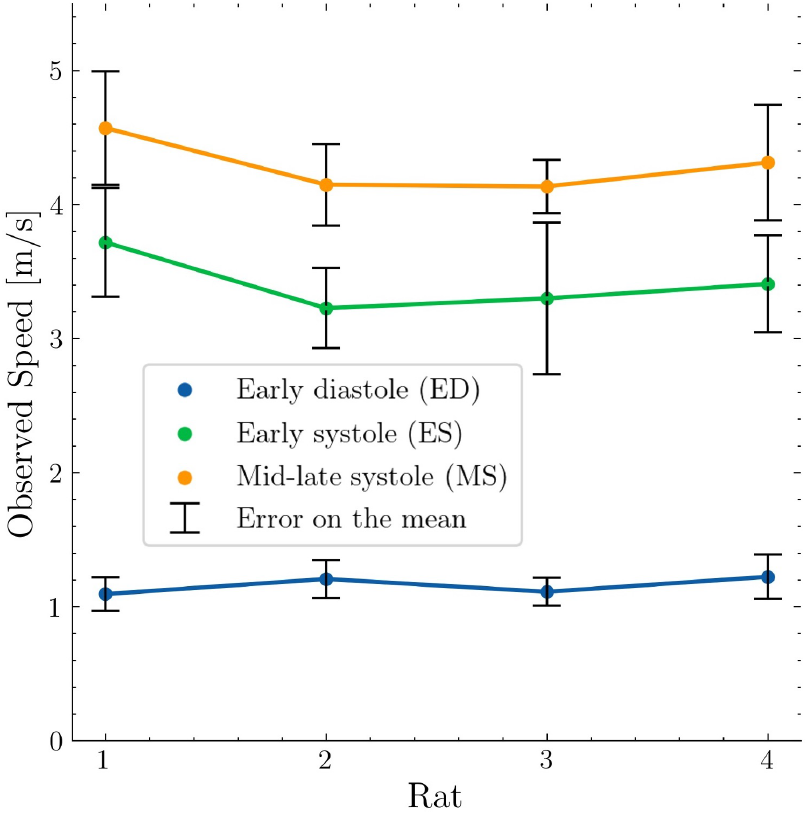
Mean apparent speed values and standard error on the mean for the three probed cardiac cycle timepoints. Aggregated data is shown grouped by animal.

MEM parameter estimations are summarized in table 1. Population mean MS speed value was estimated through the model intercept as 4.288 ± 0.088 m/s. The difference in average observed velocity between MS and ES and MS and ED were found to be significant (p-value < 0.001) estimated via the fixed effect slope parameters as 0.859 ± 0.043 and 3.135 ± 0.091 m/s respectively.

**Table 1.** Mixed effects model statistics estimations. MS was used as the reference cardiac phase (*β*_0_, *γ*_0_).

|  | Mean | Standard Error | p-value | 95% CI |  |
| --- | --- | --- | --- | --- | --- |
| $\beta_0$ | 4.288 | 0.088 | $< 0.001$ | 4.115 | 4.460 |
| $\beta^{(ED)}$ | -3.135 | 0.091 | $< 0.001$ | -3.314 | -2.956 |
| $\beta^{(ES)}$ | -0.859 | 0.043 | $< 0.001$ | -0.943 | -0.774 |
| $\text{Var}(\gamma_0)$ | 0.058 | 0.657 | | | |
| $\text{Cov}(\gamma_0, \gamma_1^{(ED)})$ | -0.058 | 0.664 | | | |
| $\text{Var}(\gamma_1^{(ED)})$ | 0.058 | 0.688 | | | |
| $\text{Cov}(\gamma_0, \gamma_1^{(ES)})$ | 0.017 | 0.263 | | | |
| $\text{Var}(\gamma_1^{(ES)})$ | 0.006 | 0.076 | | | |
| $\text{Cov}(\gamma_1^{(ED)}, \gamma_1^{(ES)})$ | -0.016 | 0.265 | | | |

Values for ED, where the heart relaxes to get filled with blood, were significantly lower than in both systole phases. MS values, where ventricular pressure is at its peak, displayed higher velocity values than ES, where the myocardium is still contracting.

From the MEM statistical moment estimations (table 1) we found a *CC*_*ED,MS*_ of −1 and a *CC*_*ES,MS*_ of 0.9113. A *CC* = −1 corresponds to perfect inversely correlation between magnitudes, while two variables with a *CC* = 1 grow or decrease coherently.

### Validation and bias correction

Table 2 displays the mean and standard error estimations for the interventricular septum thickness for each animal and each cardiac phase, as well as apparent velocity, unbiased speed, and stiffness. All three tMRE estimations of gel phantom stiffness were coherent with steady state MRE and the expected value of ∼ 1 kPa (table 2). We found corrected speed and stiffness values that are physiologically implausible for ES and MS (data not shown). We determined Mo et al. empirical law as not applicable outside the region of data support in the original work^23^ (figure 5).

**Table 2.** Experimental estimations of the mean and standard error for septum thickness (*H*), apparent shear wave speed (*C*), its corrected value (*C*_*s*_) and the corresponding shear wave modulus (*G*^*′*^). Values have been rounded to three significant figures. Numbers in the “animal” column refer to each of the 4 wistar rat subjects except for phantom experiments where these correspond to the three tMRE repetitions and the steady state MRE estimations. †: Mo et al. correction was determined as non-applicable for ES and MS (data not shown).

| Cardiac phase | Animal | $H$ (mm) | $C$ (m/s) | $C_s$ (m/s) | $G'$ (kPa) |
| --- | --- | --- | --- | --- | --- |
| Early diastole | 1 | $1.35 \pm 0.07$ | $1.09 \pm 0.13$ | $2.90 \pm 0.50$ | $8.89 \pm 3.61$ |
| | 2 | $1.67 \pm 0.05$ | $1.21 \pm 0.14$ | $2.93 \pm 0.61$ | $9.05 \pm 3.74$ |
| | 3 | $2.01 \pm 0.05$ | $1.11 \pm 0.10$ | $2.22 \pm 0.35$ | $5.20 \pm 1.65$ |
| | 4 | $1.31 \pm 0.06$ | $1.21 \pm 0.17$ | $3.63 \pm 0.89$ | $13.9 \pm 6.79$ |
| Early systole | 1 | $2.51 \pm 0.07$ | $3.72 \pm 0.41$ | † | † |
| | 2 | $2.77 \pm 0.07$ | $3.23 \pm 0.30$ | † | † |
| | 3 | $2.38 \pm 0.09$ | $3.30 \pm 0.57$ | † | † |
| | 4 | $2.08 \pm 0.05$ | $3.41 \pm 0.36$ | † | † |
| Mid-late systole | 1 | $1.61 \pm 0.07$ | $4.57 \pm 0.42$ | † | † |
| | 2 | $2.21 \pm 0.06$ | $4.15 \pm 0.30$ | † | † |
| | 3 | $2.15 \pm 0.08$ | $4.13 \pm 0.20$ | † | † |
| | 4 | $1.88 \pm 0.05$ | $4.31 \pm 0.43$ | † | † |
| Phantom | tMRE I | - | - | $1.09 \pm 0.12$ | $1.19 \pm 0.26$ |
| | tMRE II | - | - | $0.97 \pm 0.16$ | $0.94 \pm 0.31$ |
| | tMRE III | - | - | $1.03 \pm 0.12$ | $1.06 \pm 0.25$ |
| | steady state MRE | - | - | $1.02 \pm 0.17$ | $1.04 \pm 0.35$ |

**Figure 5.**
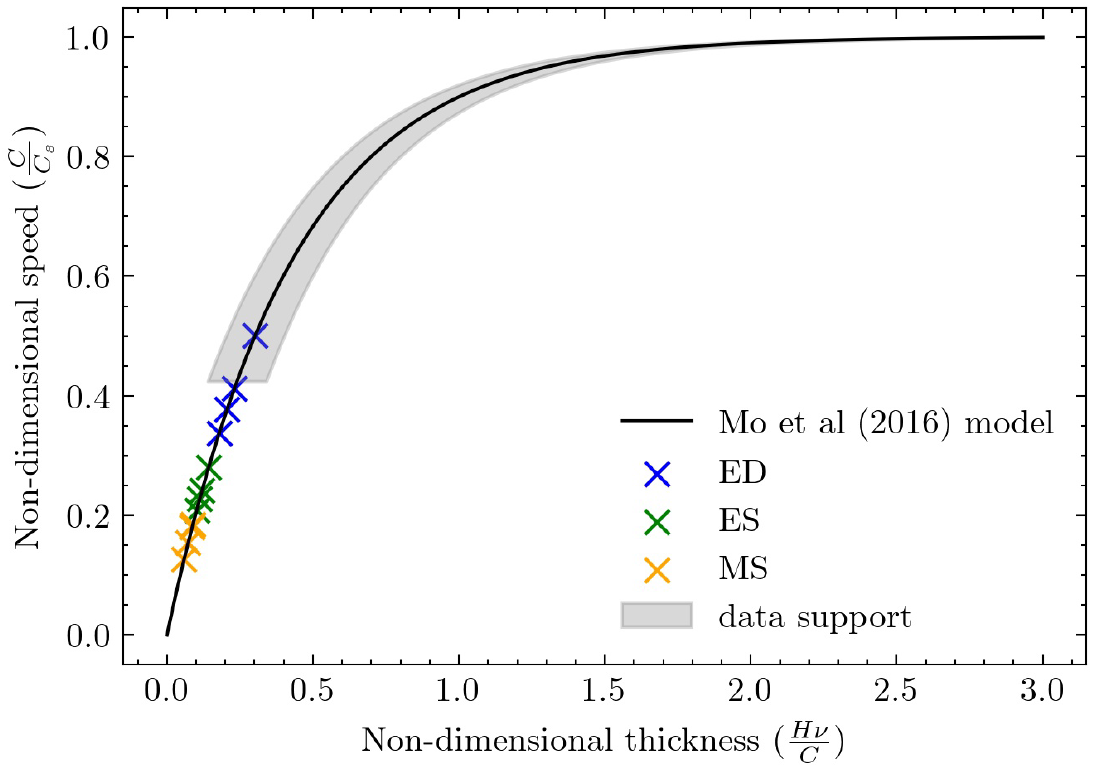
Reproduction of figure 7 in Mo et al.^23^. The dimensionless representation of the geometrical correction model is overlaid to our data. Shaded area represents the region where the authors obtained empirical data to infer the correction law.

## Discussion

In this paper we presented tMRE as a new non-invasive technique to measure shear wave speed as thereof travel through heart muscle in living animals. Shear wave speed has a know direct relationship with tissue stiffness and hence tMRE measurements have the potential of local biomarker for myocardium health status. As a proof-of-concept we quantified shear wave velocity in early diastole, early and mid-late systole in 4 different rat specimens. Our data acquisition and quantification pipeline was repeatable and reproducible across intra-specimen experiments and healthy animals. tMRE measurements had high face validity showing a clear distinction and non-overlapping of speed values at all three phases consistent with cardiac physiology. The proposed tMRE methodology was validated in phantom experiments where the material stiffness was known.

The key component of tMRE, and hence our main contribution, is the elegant overcoming of hardware constraints in capturing shear wave perturbations. tMRE is able to capture shear wave perturbations with high temporal resolution that allows for accurate downstream speed quantification. This is achieved by reorganizing repeated motion-sensitive imaging acquisitions with different time delays synchronized by the animal ECG R-peak. This DAQ strategy is highly sensitive to movement hence we equipped tMRE with various normalization and denoising tools to maximize the quality of the data. One main advantage of tMRE over previously published work is its flexibility which allows to probe cardiac biomechanics *in vivo* at any timepoint in the cardiac cycle.

Apparent speed values suffer from thin plate bias of variable magnitude depending on the septum thickness in each cardiac phase. Factors such as age, congenital anatomical differences and genetics also affect heart geometry across individuals. We used Mo et al.^23^ correction law on the region supported by simulation data based on non-dimensional speed and thickness. We found that this model was only applicable to our *in vivo* measurements for ED. Reported ED shear modulus are consistent with value ranges previously reported^25^ for adult rats myocardium assuming Poisson ratios between 0.45 and 0.5. However, measurements for ES and MS fell outside the region of support and showed disproportionally high stiffness for biological soft tissue. Our findings suggest that an ad-hoc correction law is needed in order to accurately remove the heart thin-plate bias and reveal the true tissue mechanical properties. Future work will run in this direction exploring the possibility make to make computational simulations for various heart geometries.

Nonetheless, applying Mo et al.^23^ law to ED data allowed us to validate tMRE measurements for both systolic phases. Thin-plate geometry bias is known to be an underestimation bias, meaning that the measured apparent value is lower bound for the true velocities^14,15^. We found that true speed for ED was lower than biased velocity values in ES and MS. Therefore, in the application of the appropriate correction law for the systolic observations, and recalling that the transformation from speed to modulus is monotonic, the stiffness ordering among cardiac phases will remain. This validates tMRE sensitivity to differences in myocardium stiffness across different cardiac cycle phases.

Furthermore, were also able to unveil useful patterns from the apparent speed measurements. We found two interesting signals analyzing the covariance of the data through MEM CCs. First, animals with a larger MS biased speed tend to have a lower ED velocity values, which could potentially reflect that hearts that cannot fully contract, cannot completely relax either. This may indicate that the velocity span across the cardiac cycle may be a biomarker for overall heart function. Secondly, individuals with high ES speed values tend to show high MS velocities as well. This may be expected in healthy individuals but may not hold in certain systolic conditions such as obstructive hypertrophic cardiomyopathy with dynamic outflow obstruction and/or individuals under certain treatments like beta-blockers that reduce contraction strength.

Leveraging tMRE could enable a plethora of applications in the *in vivo* imaging realm such as early diagnosis of CVDs (heart failure, myocarditis, etc), treatment monitoring and evaluation of new drugs.

## Contributions

In this work we made the following contributions:

- We presented cardiac tMRE, a novel non-invassive imaging technique aimed to estimate the stiffness of myocardium tissue *in vivo* at any desired cardiac cycle timepoint.
- We used tMRE to measure shear wave phase velocity for three different cardiac phases: early systole, mid-late systole, and early diastole.
- We found significant systematic differences in apparent speed among all cardiac phases consitent with the heart dynamic physiology.
- We provided with absolute shear modulus (stiffness) values at ED utilizing a speed correction law from the literature.

## Acknowledgements

This work was performed in collaboration with the the IAME group (UMR1137) and the Fédération de Recherche en Imagerie Multimodale facility (FRIM, UMS34) hosted at University Paris Cité and member of the France Life Imaging (grant ANR-11-INBS-0006) and IDEX Imagerie Du Vivant networks.

## Author contributions statement

MBM conceived the experiments. MBM conducted data collection with aid from GA and GR. MBM performed data analysis and modeling. MBM and JM interpreted the results. MBM and JM wrote this manuscript with critical review from all authors.

## Data and code availability

Data and code are available upon reasonable request to the corresponding author.

## Additional information

The authors manifest no competing interest.

## References

1. Voorhees, A. P. & Han, H. C. Biomechanics of cardiac function. Compr. Physiol. 5, 1623–1644, DOI: 10.1002/cphy.c140070 (2015).

2. Knight, W. E. et al. Ex vivo Methods for Measuring Cardiac Muscle Mechanical Properties. Front. Physiol. 11, 616996, DOI: 10.3389/FPHYS.2020.616996 (2020).

3. Holzapfel, G. A. Computational Biomechanics of Soft Biological Tissues: Arterial Walls, Hearts Walls, and Ligaments. Encycl. Comput. Mech. Second. Ed. 1–47, DOI: 10.1002/9781119176817.ECM2041 (2017).

4. Gazewood, J. & Turner, P. L. Heart Failure with Preserved Ejection Fraction: Diagnosis and Management. Am. family physician (2017).

5. From, A. M. & Borlaug, B. A. Heart failure with preserved ejection fraction: Pathophysiology and emerging therapies. Cardiovasc. Ther. 29, DOI: 10.1111/J.1755-5922.2010.00133.X, (2011).

6. Kovács, S. J. Diastolic function in heart failure. Clin. Medicine Insights: Cardiol. 9, 49–55, DOI: 10.4137/CMC.S18743 (2015).

7. Borlaug, B. A. Evaluation and management of heart failure with preserved ejection fraction, DOI: 10.1038/s41569-020-0363-2 (2020).

8. Emig, R. et al. Passive myocardial mechanical properties: meaning, measurement, models. Biophys. Rev. 13, 587, DOI: 10.1007/S12551-021-00838-1 (2021).

9. Nagueh, S. F. Left Ventricular Diastolic Function: Understanding Pathophysiology, Diagnosis, and Prognosis With Echocardiography. JACC: Cardiovasc. Imaging 13, 228–244, DOI: 10.1016/J.JCMG.2018.10.038 (2020).

10. O, F. Zur Dynamik des Herzmuskels. Zeitschr. Biol. 32, 370–437 (1895).

11. Sagawa, K. The ventricular pressure-volume diagram revisited. Circ. Res. 43, 677–687, DOI: 10.1161/01.RES.43.5.677/ASSET/89017E2F-059C-4E82-B7D2-DFD3E2EE17CF/ASSETS/01.RES.43.5.677.FP.PNG (1978).

12. Sinkus, R. et al. High-resolution tensor MR elastography for breast tumour detection. Phys. medicine biology 45, 1649–1664, DOI: 10.1088/0031-9155/45/6/317 (2000).

13. Khan, S., Fakhouri, F., Majeed, W. & Kolipaka, A. Cardiovascular magnetic resonance elastography: A review, DOI: 10.1002/nbm.3853 (2018).

14. Sinkus, R., Daire, J.-L., Vilgrain, V. & E. Van Beers, B. Elasticity Imaging via MRI: Basics, Overcoming the Waveguide Limit, and Clinical Liver Results. Curr. Med. Imaging Rev. 8, 56–63, DOI: 10.2174/157340512799220544 (2012).

15. Troelstra, M. A. et al. Shear wave cardiovascular MR elastography using intrinsic cardiac motion for transducer-free non-invasive evaluation of myocardial shear wave velocity. Sci. Reports 11, 1403, DOI: 10.1038/s41598-020-79231-z (2021).

16. Liu, Y., Royston, T. J., Klatt, D. & Lewandowski, E. D. Cardiac MR elastography of the mouse: Initial results. Magn. resonance medicine 76, 1879–1886, DOI: 10.1002/MRM.26030 (2016).

17. Smith, L., Skulberg, V., Zhang, L., Sjaastad, I. & Espe, E. The effects of geometry on stiffness measurements in high-field magnetic resonance elastography: A study on rodent cardiac phantoms. J. Mech. Behav. Biomed. Mater. 133, 105302, DOI: 10.1016/J.JMBBM.2022.105302 (2022).

18. Savitzky, A. & Golay, M. J. Smoothing and Differentiation of Data by Simplified Least Squares Procedures. Anal. Chem. 36, 1627–1639, DOI: 10.1021/AC60214A047/ASSET/AC60214A047.FP.PNG_V03 (1964).

19. Baayen, R. H., Davidson, D. J. & Bates, D. M. Mixed-effects modeling with crossed random effects for subjects and items. J. Mem. Lang. 59, 390–412, DOI: 10.1016/J.JML.2007.12.005 (2008).

20. Garteiser, P. et al. Rapid acquisition of multifrequency, multislice and multidirectional MR elastography data with a fractionally encoded gradient echo sequence. NMR biomedicine 26, 1326–1335, DOI: 10.1002/NBM.2958 (2013).

21. Sinkus, R. et al. Viscoelastic shear properties of in vivo breast lesions measured by MR elastography. Magn. resonance imaging 23, 159–165, DOI: 10.1016/J.MRI.2004.11.060 (2005).

22. Erich Leo Lehmann. Elements of Large-Sample Theory. Elem. Large-Sample Theory DOI: 10.1007/B98855 (1999).

23. Mo, J. et al. Bias of shear wave elasticity measurements in thin layer samples and a simple correction strategy. SpringerPlus 5, DOI: 10.1186/S40064-016-2937-3 (2016).

24. Schneider, C. A., Rasband, W. S. & Eliceiri, K. W. NIH Image to ImageJ: 25 years of image analysis. Nat. Methods 2012 9:7 9, 671–675, DOI: 10.1038/nmeth.2089 (2012).

25. Bhana, B. et al. Influence of substrate stiffness on the phenotype of heart cells. Biotechnol. bioengineering 105, 1148–1160, DOI: 10.1002/BIT.22647 (2010).

