## Supplementary Materials for "Transient Magnetic Resonance Elastography: a method to measure the mechanics of the active heart"

#### Quantitative cardiac cycle biomechanics: An MRE measurement of the active heart

##### 1 Additional figures

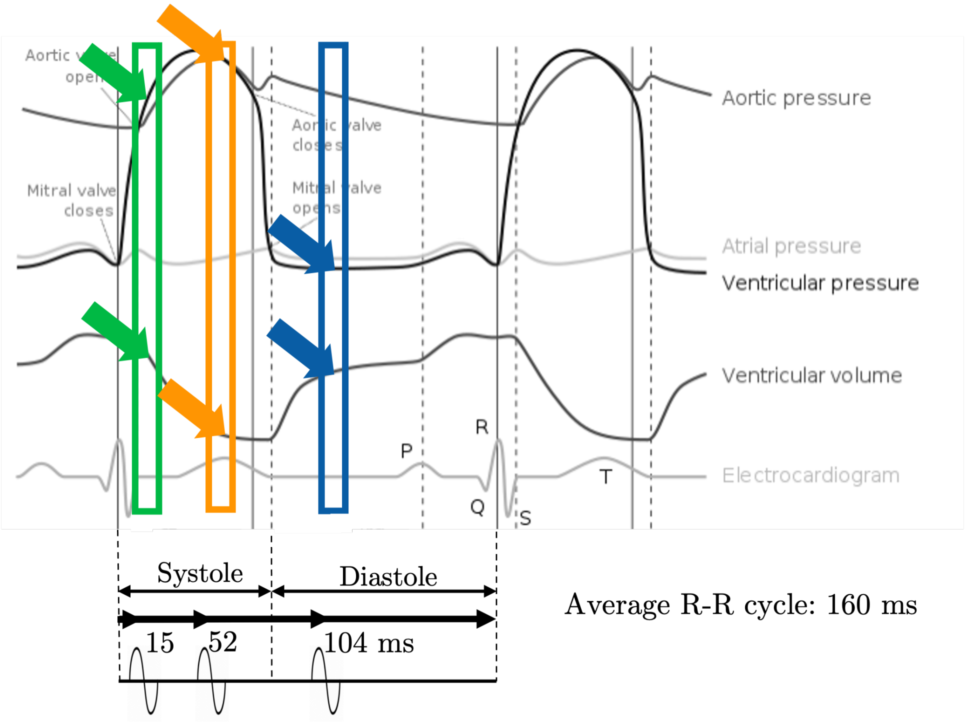


Figure S1: Wiggers diagram displaying the ventricular and atrial relative volumes and pressure at the three probed cardiac cycle timepoints.

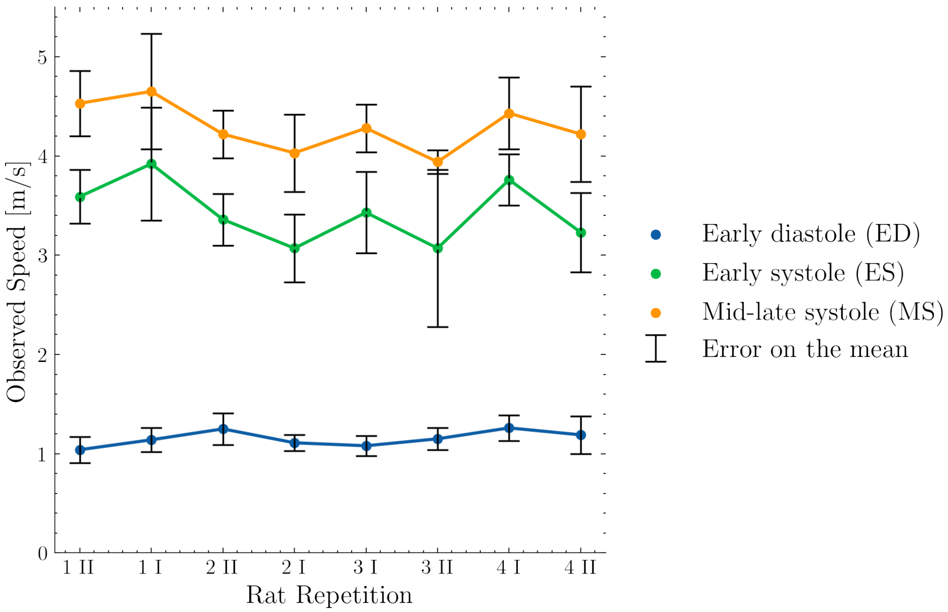


Figure S2: Mean apparent speed values and standard error on the mean for the three probed cardiac cycle timepoints for each experiment. Aggregated data is shown grouped by repetition. Arabic numbers refer to the specimen while roman figures indicate repetition.

### 2 Additional methods

#### 2.1 Mixed effects modelling

We fitted a mixed effects model (MEM) to the mean apparent speed data across specimens and repetitions. The interesting characteristic of MEMs is that they allow the individual estimation of both the true systematic differences between cardiac phases and the experimental random variation (random effects).

In order to fulfill model assumptions we assumed independence among experiments (rat and repetition) observations, errors and random variation effects. We defined the MEM as

$$y_i^{(j)} = x_i \boldsymbol{\beta} + z_i \boldsymbol{\gamma_i} + \boldsymbol{\gamma_{i,0}} + \epsilon_i^{(j)}, \quad (1)$$

with

$$\epsilon_i \sim \mathcal{N}(0, \sigma^2) \quad (2)$$

being $\sigma^2$ unknown, and

$$\gamma_0 \sim \mathcal{N}(0, Var(\gamma_0)) \quad ; \quad \gamma_1^{(j)} \sim \mathcal{N}(0, Var(\gamma_1^{(j)})) \quad (3)$$

We define $y_i^{(j)}$ as the observed speed for the $i^{th}$ experiment and $j^{th}$ cardiac phase;$x_i$ and $z_i$ represent the fixed and random effects design matrices columns for the $i^{th}$ experiment for the three cardiac phases, respectively; $\boldsymbol{\beta}$ estimates the fixed effects coefficient vector, hence it encodes the speed differences between cardiac phases plus an intercept that is interpreted as the mean speed corresponding to the reference cardiac phase, Mid-late systole (MS) in our setting; contrarily, $\boldsymbol{\gamma_{i,1}}$ allows inference on the random effects through a random coefficient vector whose $j^{th}$ elements are drawn from the normal distributions shown above at each $i^{th}$ instance. In plain terms, it models the random variation on pairwise speed differences across cardiac phases; $\epsilon_i^{(j)}$ denotes the error in each measurement.

The calculation of the correlation coefficients (CCs) for early systole (ES) and early diastole (ED) enabled us to quantify trends stemming from experimental variability in the observed speed at each cardiac phase in our sample. We calculate the CC between the $j^{th}$ cardiac phase and MS (reference) as

$$CC_{j,MS} = \frac{Cov(\gamma_0, \gamma_1^{(j)})}{\sqrt{Var(\gamma_0) Var(\gamma_1^{(j)})}}. \quad (4)$$

We used the python package *statsmodels* for the implementation of our MEM.

#### 2.2 Standard error estimation via the delta method

In summary, the delta method states that if a random variable $X_n$ satisfies

$$\sqrt{n} \times [X_n - \mu] \xrightarrow{D} \mathcal{N}(0, \sigma^2), \quad (5)$$

with population mean $\mu$, sample size $n$ and population variance $\sigma^2$, it follows that

$$\sqrt{n} \times [g(X_n) - g(\mu)] \xrightarrow{D} \mathcal{N}(0, \sigma^2 \times [g'(\mu)]^2) \,, \tag{6}$$

for a differentiable function $g(.)$ that fulfills $g'(\mu) \neq 0$.

We can use this result to approximate the uncertainty of an estimator for a non-linearly transformed variable. Therefore, for each *in vivo* experiment shear wave velocity ($C$) we make the reasonable assumption that

$$C \sim \mathcal{N}(\mu_C, \sigma_C^2) \,, \tag{7}$$

which follows the law of big numbers and satisfies equation 5.

Thus, we can apply the delta method to find the variance of the corrected shear wave speed ($C_s$) and of shear modulus ($G'$):

- In the first case we consider the non-linear transformation

  $$C_s = g(C) = C/(1 - e^{-2.3\times\frac{H\times\nu}{C}}) \,, \tag{8}$$

  and approximate the standard error on the mean for $C_s$ via the delta method as

  $$SE_{C_s} = \frac{\hat{\sigma_C} \times g'(\bar{C})}{\sqrt{n}} \,, \tag{9}$$

  where the population statistics $\mu_C$ and $\sigma_C$ are approximated with the sample estimates $\bar{C}$ and $\hat{\sigma_C}$, and

  $$g'(\bar{C}) = \frac{1 + e^{-2.3\times\frac{H\nu}{C}} \times (2.3 \times \frac{H\nu}{C} - 1)}{(1 - e^{-2.3\times\frac{H\nu}{C}})^2} \,. \tag{10}$$

- The second non-linear relationship corresponds to combining equation 8 with

  $$G' = \rho \times C_s^2 \,, \tag{11}$$

  to obtain

  $$G' = \tilde{g}(C) = \rho \times C^2/(1 - e^{-2.3\times\frac{H\nu}{C}})^2 \,. \tag{12}$$

  The parameter $\rho$ corresponds to the myocardium density which has been estimated as 1055 kg/m$^3$ [1].

  Following the same procedure as before we can approximate the standard error on the mean for $G'$ as

  $$SE_{G'} = \frac{\hat{\sigma_C} \times \tilde{g}'(\bar{C})}{\sqrt{n}} \,, \tag{13}$$

  with

  $$\tilde{g}'(\bar{C}) = 2\rho C \left[ \frac{1 + e^{-2.3\times\frac{H\nu}{C}} \times (2.3 \times \frac{H\nu}{C} - 1)}{(1 - e^{-2.3\times\frac{H\nu}{C}})^3} \right] \,. \tag{14}$$

For the phantom stiffness estimation we can apply a similar rationale and water density ($\rho = 1000$ kg/m$^3$). In this case we measure $C_s$ directly thus we assume:

$$C_s \sim \mathcal{N}(\mu_{C_s}, \sigma^2_{C_s}) \ . \tag{15}$$

We then apply the delta method as before to obtain standard errors for $G'$ using equation 13 and

$$\tilde{g}'(C_s) = 2\rho C_s \ . \tag{16}$$

### References


[1] Bishwas Chamling, Michael Bietenbeck, Stefanos Drakos, Dennis Korthals, Volker Vehof, Philipp Stalling, Claudia Meier, and Ali Yilmaz. A compartment-based myocardial density approach helps to solve the native T1 vs. ECV paradox in cardiac amyloidosis. *Scientific Reports 2022 12:1*, 12(1):1–9, 12 2022. URL: https://www.nature.com/articles/s41598-022-26216-9, doi:10.1038/s41598-022-26216-9.
